# *In Vivo* Simultaneous Nonlinear Absorption Raman and Fluorescence (SNARF) Imaging of Mouse Brain Cortical Structures

**DOI:** 10.1101/2021.06.26.450059

**Authors:** Andrew T. Francis, Bryce Manifold, Elena C. Thomas, Ruoqian Hu, Andrew H. Hill, Shuaiqian Men, Dan Fu

**Affiliations:** Department of Chemistry, University of Washington, Seattle, Washington 98195, United States

## Abstract

Two-photon excited fluorescence (TPEF) microscopy is a widely used optical imaging technique that has revolutionized neurophotonics through a diverse palette of dyes, specialized transgenic models, easy implementation, and straightforward data analysis. However, *in vivo* TPEF imaging is often limited in the number of contrasts available to distinguish different cells, structures, or functions. We propose using two label-free multiphoton microscopy techniques – stimulated Raman scattering (SRS) microscopy and transient absorption microscopy (TAM) – as complementary and orthogonal imaging modalities to TPEF for *in vivo* brain imaging. In this study, we construct a simultaneous nonlinear absorption, Raman, and fluorescence (SNARF) microscope and image several cortical structures up to 250-300 μm below the pial surface, the highest reported *in vivo* imaging depth for SRS or TAM. We further demonstrate the capabilities of our SNARF microscope through the quantification of age-dependent myelination, hemodynamics, vessel structure, cell density, and cell identity *in vivo.* Using machine learning, we report the use of label-free SRS and TAM features to predict capillary-lining cell identities with 90% accuracy. The SNARF microscope and methodology outlined herein provide a powerful platform to study several research topics, including neurovascular coupling, blood-brain barrier, neuronal and axonal degeneration in aging, and neurodegenerative diseases.

## Introduction

The application of laser scanning microscopy to neuroscience has revolutionized our understanding of brain structure, development, and function under physiological and pathological conditions. Two-photon excited fluorescence (TPEF) has emerged as a leading brain imaging technique due to its high spatial resolution, intrinsic optical sectioning, and technical simplicity.^1–3^ Advancements in fluorescent transgenic animals and highly specific fluorescent dyes have empowered researchers to precisely label cell types and structures *in vivo*. However, despite tremendous technical development over the years, TPEF suffers from multiple limitations: 1) TPEF emission spectra are typically broad (~50 nm), which limits the number of separable fluorescent contrasts, and thus, information that can be collected from one animal, 2) generating animals with multiple transgenes can be challenging, time-consuming, and expensive^4^ and 3) application of fluorescent dyes – topically or intravenously – is invasive which can alter the animal’s physiology.^5^

To circumvent these limitations, orthogonal label-free multiphoton imaging techniques can generate additional image contrasts. Third harmonic generation (THG) is one such method that provides characterization of myelination and cell density in the brain.^6,7^ While robust, THG does not provide molecular information. Instead, researchers have explored the use of coherent Raman scattering (CRS) microscopy for label-free chemical imaging. Coherent anti-Stokes Raman scattering (CARS) and stimulated Raman scattering (SRS) – two CRS techniques – have shown tremendous capability in the chemical mapping of endogenous lipids, proteins, and other biomolecules in animal and human tissues.^8–13^ SRS has several advantages over CARS, including a linear dependence between signal size and chemical concentration and the absence of a non-resonant background, making SRS the more widely employed of the two CRS techniques. One emerging application of SRS microscopy is the development of label-free histopathology, which achieves comparable diagnostic insight as traditional staining methods (e.g., H&E).^14,15^ Despite its capabilities, reports of *in vivo* brain imaging with SRS remains scarce largely due to poor penetration depth. Ji *et al.* reported an SRS imaging depth of ~100 μm when demonstrating *in vivo* pathological differentiation between tumor and nonneoplastic tissue in a xenografted mouse brain model.^16^ To address this issue, our group and others have worked to increase signal sizes and thus penetration depth, via pulse optimization^17^, wavelength optimization^18^, aberration correction^19^, tissue clearing^20,21^, and deep learning^22,23^. To date, these improvements have so far only been demonstrated *ex vivo*.

Transient absorption microscopy (TAM) is another label-free chemical imaging technique that is potentially useful for *in vivo* brain imaging. Recent reports have used the intrinsic absorption of hemoglobin to quantify hemoglobin glycation,^24^ oxygenation,^25^ and concentration^26^ *ex vivo*. Importantly, imaging of hemoglobin allows direct visualization of blood flow and relieves the need for intravenous fluorescent dye injections into the blood compartment, which are invasive and can suffer from quick clearing times.^27^ However, there has been no attempt in *in vivo* TAM imaging of the brain.

In this study, we constructed a <u>s</u>imultaneous <u>n</u>onlinear <u>a</u>bsorption, <u>R</u>aman, and <u>f</u>luorescence (SNARF) microscope for *in vivo* mouse brain imaging. The SNARF microscope combined TAM, SRS, and TPEF to simultaneously interrogate proteins, lipids, hemoglobin, and fluorophores. Through pulse optimization and denoising, we performed the deepest *in vivo* SRS imaging of brain tissue at ~300 μm and the first *in vivo* TAM imaging of the brain. We employed SNARF microscopy to characterize, quantify, and compare the differences in cell density, myelination, and hemodynamics between three groups of mice – young (P30-P50), middle-aged (P200-P250), and old (P630-P635). Using TPEF of a Tie2-GFP transgenic mouse stained with Neurotrace 500/525 as ground truth, we built a Random Forest classifier for label-free identification of two classes of capillary-lining cells – endothelial cells and pericytes – based on morphology and relation to neighboring capillaries with 90 ± 2% accuracy. SNARF microscopy builds upon the powerful capabilities of TPEF microscopy by adding orthogonal and complementary chemical contrasts that would otherwise require additional fluorescent labeling. We believe that a more comprehensive and detailed *in vivo* imaging platform will benefit several neuroscience research areas, including neurovascular coupling, blood-brain barrier, cerebral amyloid plaque, myelin degeneration and regeneration, and neurodegeneration.

## Methods

### SNARF microscope system

The SNARF microscope used for imaging, shown in Figure 1A, was built upon a previously described SRS microscope system.^28^ The laser system starts with a femtosecond dual-beam laser (Spectra-Physics, Insight DS+) outputting two beams with an 80 MHz (f_0_) repetition rate. The two outputs are a fixed beam at 1040 nm, used as the pump, and a tunable beam ranging from 680 to 1300 nm, used as the probe. The pump beam (fixed at 1040 nm) was amplitude modulated at 20 MHz (f_0_/4) by an electro-optical modulator (EOM1, θ_1_) and a polarizing beam splitter (PBS). The amplitude modulated pulse train was followed by another polarization modulation at 20 MHz with a second EOM (EOM2, θ_2_). The second EOM is operated 90 degrees phase shifted from the first EOM (i.e., θ_2_ − θ_1_ = 90°), resulting in two orthogonal phase pulse trains (s and p polarization). From here, the beam was passed through a 20 mm of birefringent quartz crystal (BRC) (Union Optic, BIF-Quartz) before traversing a half waveplate (HWP) and PBS to combine both beams into a single polarization. This procedure generates two orthogonally modulated 20 MHz pump pulse trains with a 90-degree phase shift and a fixed time delay, allowing simultaneous acquisition of two-channel SRS and TAM signals at two different time delays.^26,28^

**Figure 1.**
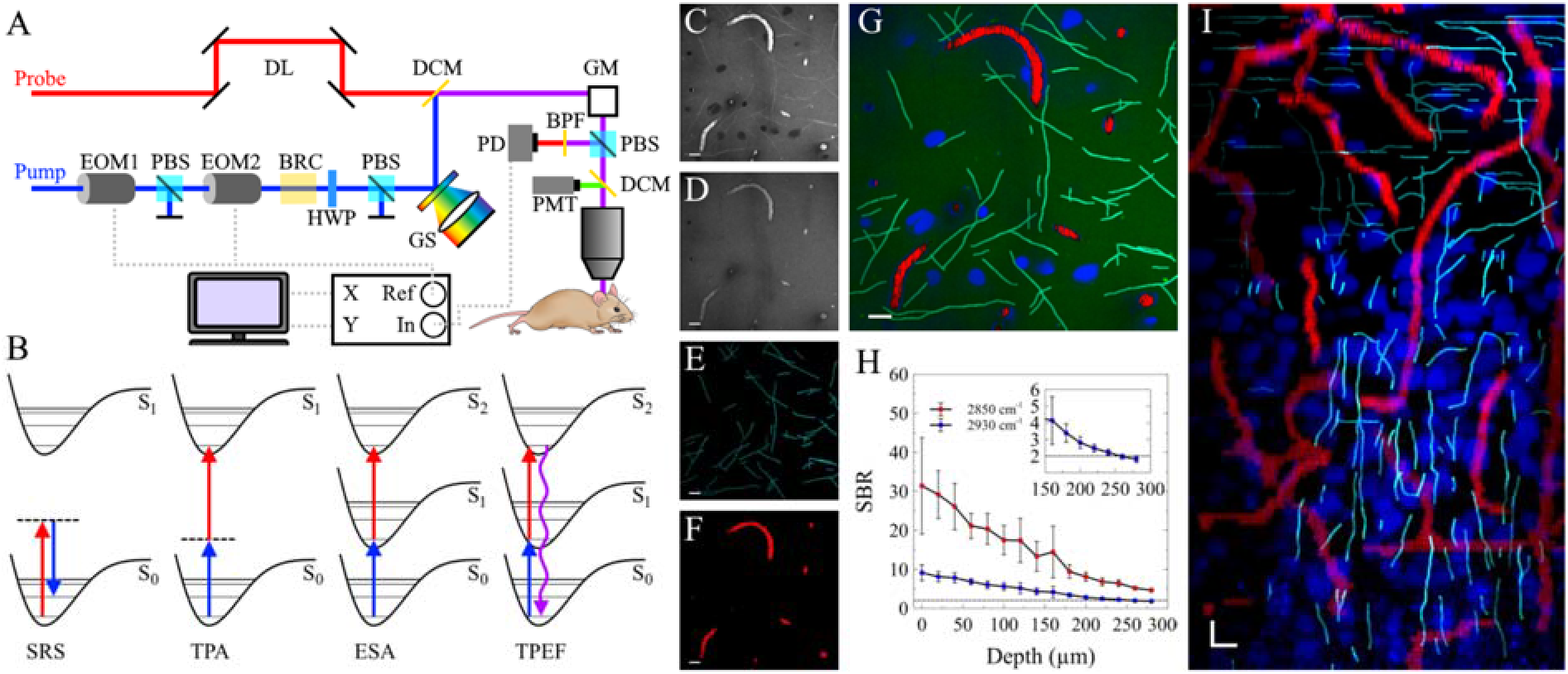
Construction of the SNARF microscope. A) Experimental set up. DL: delay line, DCM: dichroic mirror, GM: galvanomirrors, EOM: electro-optic modulator, PBS: polarizing beam splitter, BRC: birefringent crystal, HWP: half-wave plate, GS: grating-based pulse stretcher, PD: photodiode, PMT: photomultiplier tube. B) Energy level diagrams of the multiphoton techniques employed. C) and D) SRS imaging at 2850 cm^−1^ and 2930 cm^−1^, respectively. E) Extracted myelin signal. F) Extracted blood vessels. G) Composite image of lipids (green), proteins (blue), myelin (cyan), and blood vessels (red) after signal extraction and spectral unmixing. H) Signal-to-background ratio (SBR) of each wavenumber as a function of depth. I) Simulated B-slice. Scale bars: 10 μm.

Both pump pulses and the probe pulse were sent through 24 cm of highly dispersive glass rod (H-ZF52A) before being spatially overlapped using a dichroic mirror (DCM) and directed through a scanning microscope and 25X objective (Olympus XLPLN25XWMP2 NA = 1.05). Precise temporal overlap was achieved by delaying the probe pulse via a delay line (DL). Fluorescence signal was collected using a long pass DCM and photomultiplier tube (PMT). SRS and TAM signals were collected in epi-mode using a PBS and photodiode (PD). The PD signal was detected by a dual-phase lock-in amplifier (Zurich Instrument HF2LI) with orthogonal outputs after filtering out the two pump beams using two short pass filters (Thorlabs FESH1000). Two images were acquired simultaneously at a frame rate of 1 frame/sec with an image size of 512×512 pixels. The photophysical processes probed with the SNARF microscope are shown in Figure 1B.

### In vivo mouse brain imaging

All experimental animal procedures were approved by the Institute of Animal Care and Use Committee (IACUC) of the University of Washington (protocol # 4395−01). Cranial window surgery was performed on female C57BL/6 mice ranging from P37 to P632 and female Tie2-GFP mice ranging from P31-P92. All mice were purchased from Jackson Laboratory. Briefly, mice were anesthetized using 4% isoflurane (1 L/min O_2_) and a ~3 mm craniotomy was performed using a high-speed surgical hand drill. Next, fluorescent dyes – Neurotrace 500/525 or sulforhodamine 101 (SR101) – were topically applied to the exposed brain and a 5 mm coverslip was placed over the craniotomy. Neurotrace 500/525 and SR101 were applied for 5 and 10 min, respectively, as described previously.^5,29^ Before sealing the window with dental cement (Parkell), the window was filled with modified artificial cerebrospinal fluid to mitigate tissue movement during imaging.^30^

### Machine learning metrics and Random Forest (RF) algorithm

The 3D morphology of each cell was calculated using Aivia. The relationship between each cell and neighboring capillary was calculated using Matlab. The distance of each cell to the neighboring capillary was calculated from the 3D center of mass of the cell to the closest edge of a capillary. The direction of each cell was calculated by first segmenting a 5 μm cube centered around the 3D center of mass of the cell. The cell and neighboring capillary contained within the segmented cube were then each modeled as an ellipsoid. The direction was calculated by measuring the angle between the primary axis of the cell ellipsoid and the capillary ellipsoid. A Random Forest (RF) algorithm was employed to distinguish between morphological and locational features of multiple cell classes. We use a 20-tree RF algorithm with a 70/30 training/testing separation. The algorithm was trained and tested using Matlab.

## Results

SRS microscopy has been widely used to image lipids and proteins in various biological systems ranging from cells to tissue. Spectral contrast between these species is achieved by probing two transitions, 2850 cm^−1^ (lipids) and 2930 cm^−1^ (lipids and proteins), in the CH stretching region.^16^ Figures 1C and 1D provide representative images (~140×140 μm) from a P246 mouse at these two transitions that demonstrate clear structural features highlighted by the two transitions. Specifically, Figure 1C shows dark circular features representing cell nuclei and bright thin features representing lipid-rich myelin sheaths. Myelin sheaths are multilayer lipid-rich membranes that wrap around axons to provide metabolic support and increase the propagation velocity of action potentials.^31^ Both CARS and SRS have been employed previously to visualize and quantify myelination *in vivo* in the peripheral nervous system, particularly the spinal cord and nerve bundle.^32,33^ The thickness of those myelin sheaths is typically much larger than those in the gray matter of the brain and, therefore, easier to image. *In vivo* CARS imaging of myelin sheath in the brain has been attempted, but the imaging depth was limited to only 30 μm.^34^ To quantify myelination in our measurements, we employed a previously demonstrated tracing program to allow us to separate and measure myelin segments directly.^35^ Figure 1E provides the resulting image of traced myelin segments. In addition to SRS, the images provided in Figures 1C and 1D capture TAM signal of hemoglobin as well. These signals can be separated based on three characteristics: signal intensity, wavenumber dependence, and morphology. TAM hemoglobin signals are stronger than SRS signals due to the larger absorption cross-section. Additionally, in contrast to SRS, TAM signal is relatively independent of the targeted wavenumber due to the broad absorption profile of hemoglobin. Thus, TAM signal has similar intensity in both Figures 1C and 1D. Lastly, tissue structure is nearly static throughout imaging, whereas blood vessels can be identified due to flowing blood cells. These differences allow us to segment blood vessels as shown in Figure 1F. The raw images were then denoised using a previously described deep learning denoising technique^22,23^, followed by spectral unmixing to obtain a two-color protein and lipid image, similar to previously published stimulated Raman histology approach.^36^ All combined in Figure 1G, these label-free contrasts provide an instantaneous snapshot of three cortical structures: cell density, myelination, and vasculature.

The mouse cortex has six layers containing unique and distinct cellular subtypes that enable dynamic brain activity.^37^ The outermost layer, layer 1 (L1), has the lowest cellular density of the cortical layers, constituting mostly non-neuronal cells, such as astrocytes.^38^ Depending on the specific cortical region, the thickness of L1 ranges from about 100 μm to 130 μm. The transition from L1 to layer 2 (L2) is marked by a sharp increase in cells, primarily neurons, at the border between the two layers. Previous demonstrations of *in vivo* SRS brain imaging were limited to ~100 μm, which would have prevented SRS imaging from observing the transition.^16^ Imaging depth was improved to 210 μm for *ex vivo* sectioned brain tissue after deep learning-based denoising.^23^ Here, we demonstrate *in vivo* imaging depths of 300 μm and ~250 μm for 2850 cm^−1^ and 2930 cm^−1^, respectively (Figure 1H, depth limit is defined at a signal to background ratio of >2), 2.5-fold higher than previous *in vivo* work. With the SNARF microscope, we observed that the number of cells doubled near 100 μm, indicating the transition from L1 to L2 (Supporting Figure 1). This trend was observed in all mice in this study (n = 10). Figure 1I shows maximum intensity XZ axis projection from the surface down to 250 μm after separating myelin and hemoglobin signals. From this cross-section, the transition from layer 1 to layer 2 can be clearly observed by the increase in cell nuclei density around 100 μm below the pial surface. Additionally, we observed the arborization of myelin sheaths towards the surface of the brain. These structures transition from predominantly vertical segments at deeper cortical layers to horizontal myelin segments close to the pial surface.

Myelin remodeling in the brain is a highly dynamic process throughout the lifetime of an animal. Under normal aging, axons continue to undergo myelination through adolescence into adulthood.^39^ However, demyelination and myelin abnormalities, which are linked to physical and cognitive decline^40^, have also been reported during normal aging.^41^ Despite the progress made in recent years, the mechanisms behind demyelination and remyelination remain elusive. Here, we demonstrate the first *in vivo* SRS quantification of myelination in the brain. We performed 3D myelin imaging of three groups of mice: young (P37, P38, P48), middle-aged (P202, P209, P244, P246), and old (P630, P631, P631). Figure 2A provides the 3D rendering of myelin structure from a P246 mouse. The 3D rendering shows the ascension and arborization of axons from layer 2/3 into layer 1. This structure is further demonstrated in the maximum intensity projections shown in Figure 2B. The cortical surface is rich in myelin with a web-like structure of overlapping and intersecting myelin segments. Deeper into the cortex, myelin penetrates down, resulting in predominantly cross-sectional views of myelin sheaths. Previous studies of myelination using other imaging techniques have shown that it is closely related to animal age.^41–45^ Figures 2C-2E provide a representative comparison in myelin density of the first 20 μm of cortex between young (P38), middle-aged (P246), and old (P631) mice. We quantified the average myelin length over the top 60 μm of cortex between the three groups (Figure 2F). In agreement with previous reports, we observe a significant increase in myelination density with age.^41^ In addition to quantifying myelin length and density, we observed two distinct myelin features: myelin gaps and myelin ballooning. Figure 2G provides a representative image of the observed myelin gap, which likely arises from Nodes of Ranvier. In Figure 2H, we fit the intensity profile of the myelin segment to determine a full width, half maximum of 1.93 μm. We repeated this procedure for 60 myelin gaps across several fields of view of a P246 mouse to produce the histogram shown in Figure 2I with an average gap length of 2.04 ± 0.76 μm. This is longer than the published length of ~1 μm for Nodes of Ranvier,^46^, suggesting that our SRS measurements include paranodal length as well.

**Figure 2.**
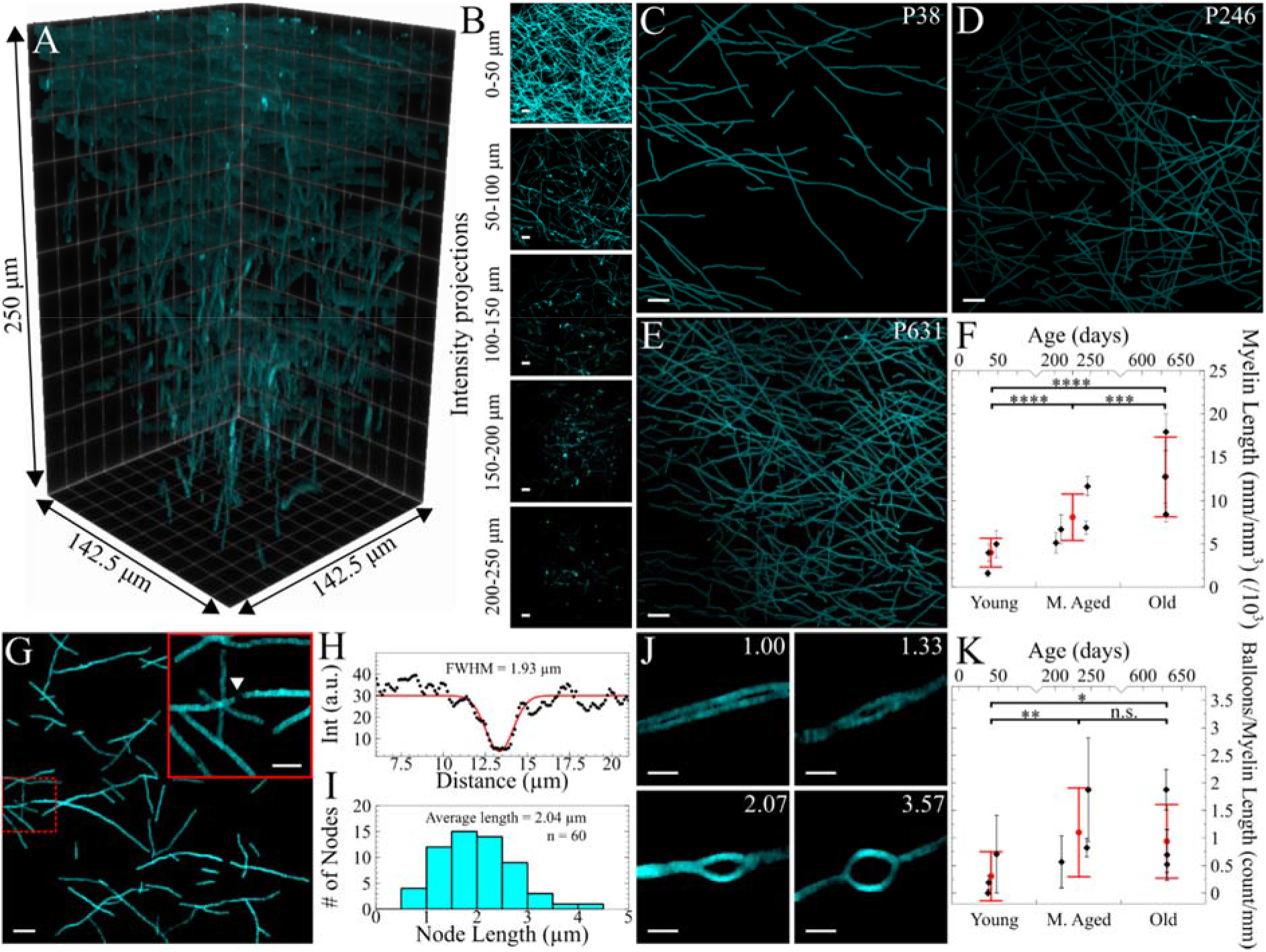
Label-free *in vivo* imaging of myelin with SNARF microscopy. A) 3D rendering of myelin. B) Intensity projections of the 3D volume shown in A) projected every 50 μm. C-E) Intensity projection of the top 20 μm of myelination of a P38, P246, and P631 mouse. F) Quantitative comparison of myelination between different aged mice. G) Suspected Node of Ranvier. H) Line profile and fitting across a myelin gap shown in G). I) Average length of suspected myelin gap. J) Quantification of myelin ballooning from not ballooning (1.00) to severe ballooning (3.57). K) Number of balloons observed per length of myelin as a function of age. Scale bars: 10 μm, G) inset scale bar: 2 μm. ****: p<0.0001, ***: p<0.001, **: p<0.01, *: p<0.1, n.s.: not significant.

We also observed myelin ballooning structures in all mouse brains imaged. We identify myelin balloons by the width ratio of the ballooned segment to the non-ballooned segment as >1.3. The ratio is reported with representative areas shown in Figure 2J. The most dramatic ballooning increased the myelin segment over 3.5 times in width. To determine if the development of myelin balloons was age-dependent, we compared the ratio of balloons to myelin length between age groups (Figure 2K). We observed an increase in balloon density between the young mouse group and the two older groups, but no significant difference between middle-aged mice and old mice. These seemingly abnormal structures could be due to brain injury.^47^ Hill *et al.* observed a similar myelin feature using SCoRe microscopy, which they classified as myelin spheroids of compact myelin due to advanced aging.^41^ From our images, axon ballooning or bulging enlarges axon diameter without apparent thickening of the myelin sheath. We also observed that most balloons enclosed a volume with similar protein and lipid concentrations to the bulk tissue. These data seem to contradict the presence of compact myelin which would give rise to strong SRS signal. In a few cases, the balloon enclosed a volume that had significantly lower signal (see Supporting Figure 2) which we believe to be a vacuole. We note that SCoRe and SRS do not provide the same contrasts: SCoRe probes light reflection due to refractive index differences and the signal is a convolution of myelin thickness, axon diameter, and surrounding material density; SRS directly images lipid density and simplifies image interpretation. Nonetheless, abnormalities in myelin structure, such as myelin swelling and ballooning, have been implicated in cognitive slowing.^48^ With its spectroscopic imaging and quantitative capability, SNARF microscopy offers a powerful alternative approach for investigating myelin degeneration and regeneration.

Next, we present label-free *in vivo* imaging of microvasculature. *In vivo* TPEF imaging of microvasculature requires an intravenous injection of a dextran-conjugated fluorophore to label the blood plasma.^30^ Here, we directly image red blood cells via the strong intrinsic absorption of hemoglobin. Supporting Figure 3 provides a representative image of a capillary labeled with FITC-dextran which demonstrates the orthogonality between TPEF and TAM for *in vivo* imaging of vasculature. Figure 3A provides a 3D rendering of the label-free TAM imaging of a ~140×140×200 μm^3^ volume of mouse brain microvasculature. Figure 3B shows the corresponding intensity projection of the volume. Recent studies have demonstrated the occurrence of stalled capillary flow in healthy and disease models, which can lead to decreased oxygen availability and neuronal injury.^49,50^ In our investigation, we observed two classes of stalled capillaries (color-coded white): transient and persistent. We identify flowing capillaries by the strong RBC contrast and stalled vessels by a noticeable decrease in protein and lipid SRS signal from the neighboring tissue. The duration of the stall (i.e., transient or persistent) was determined by imaging the same volume of tissue again after 15 minutes. Figure 3C-3D illustrates the presence of both transiently stalled vessels (green arrows, cleared before 15 min) or persistently stalled vessels (yellow arrows, remaining after 15 min). The observed prevalence of stalled vessels for each age group, ~2-5% of capillaries (see Figure 3E), is lower than reported in the literature for an acute cranial window under anesthesia.^50^ This is likely due to our inclusion of only persistently stalled vessel segments rather than all stalled vessel segments. Most stalls were cleared within a short period.^50^

**Figure 3.**
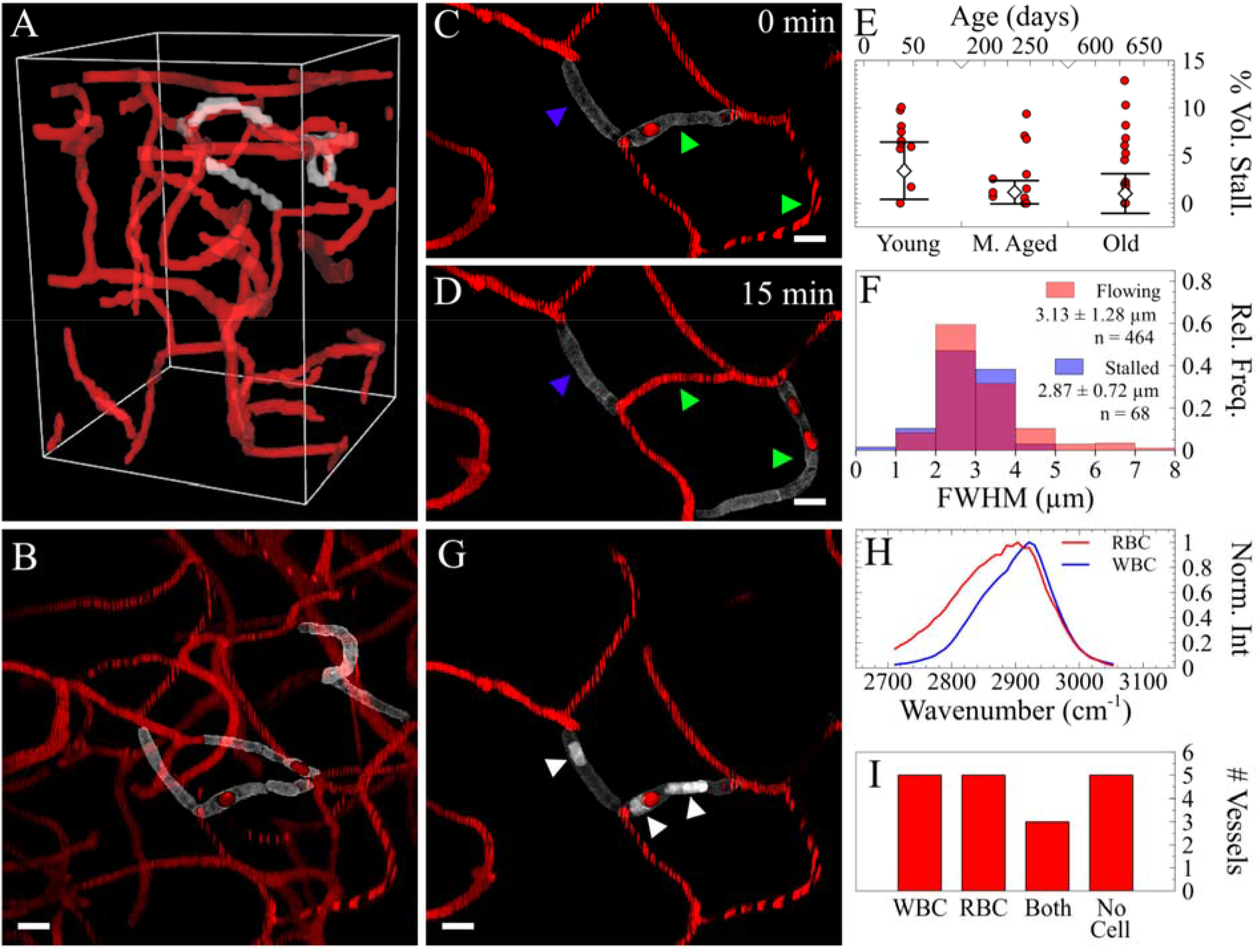
Label-free *in vivo* imaging of mouse brain microvasculature with SNARF. A) 3D rendering of mouse brain microvasculature. B) Intensity projection of flowing and stalled vessels from A). C and D) Repeated imaging of vasculature to identify stalled vessels. E) Comparison of stalled vessel volume to total vessel volume between age groups. F) Average vessel width between flowing and stalled vessels. G) SRS imaging of WBCs stalled within vasculature, indicated by white arrowheads. H) Spectral differences between RBCs and WBCs. I) Frequency of WBC, RBC, both, or no cell observed in vessels stalled for 15 min. Scale bars: 10 μm.

The cause of the stalled capillaries, and decreased cerebral blood flow, has been suggested as a cause for a variety of neurodegenerative diseases.^51^ One possible cause of stalls is a narrowing of capillaries which prevents blood cells from passing. We compared the width of 68 stalled vessel segments and 464 flowing vessel segments and found only a slight narrowing for stalled vessel width (Figure 3F). These results suggest that there is likely another cause of stalled flow. Hernandez *et al.* demonstrated that neutrophils plugging capillaries led to decreased cerebral blood flow in an Alzheimer’s Disease model.^52^ To observe those obstructions, the injection of a fluorescent dye to visualize platelets and white blood cells was necessary.^49,52,53^ Here, we leverage the label-free imaging capabilities of SRS to investigate the occurrence of white blood cells (WBCs) within the persistently stalled capillaries. The WBC nuclei, indicated by the white arrows in Figure 3G, can be separated from RBCs based on intrinsic spectral differences, as shown in Figure 3H. Stratifying the two types of blood cells allowed us to investigate the contents of stalled capillaries. We found four categories: stalled with 1) WBC(s), 2) RBC(s), 3) both WBC(s) and RBC(s), or 4) no cell. Figure 3I provides the frequency of each category over the 18 stalled capillaries identified. Only vessels that were fully contained within the imaging volume were considered. We found that ~45% (8/18 persistently stalled capillaries) contained at least one WBC. Given the low concentration of WBCs within the circulating bloodstream, these data suggest that WBCs have a higher chance of being stuck within, or plugging, a stalled capillary than RBCs. However, a larger number of stalls only have RBCs or no cells, highlighting the need to further explore the extent of stalled flow and underlying causes. It is also interesting to investigate how transient and persistent stalled flows contribute to local oxygen depletion.

Capillaries are formed by a single layer of endothelial cells. The small capillary diameter helps efficient exchange of oxygen, nutrients, and waste between the lumen of the capillary and the surrounding tissue. Dilation and contraction of capillary diameter are controlled by pericytes, cells that are embedded within the walls of capillaries.^54^ Both endothelial cells and pericytes are essential for the formation and maintenance of blood brain barrier (BBB) and regulation of immune cell entry to the central nervous system. TPEF imaging is widely used to study their functions in physiological and pathological conditions.^55,56^ Either transgenic mice or cell-specific dyes are required to image these cells. Due to the limited color channels (typically 2 to 3) that can be simultaneously acquired in *in vivo* TPEF microscopy, the number of cell types and their interactions that can be studied is restricted. Here, we perform *in vivo* SNARF imaging of Tie2-GFP mice to show multiplexing capability and develop label-free identification of capillary-lining cells. Figure 4A provides a 25 μm maximum intensity projection of SNARF imaging of Tie2-GFP in layer 1 of a Tie2-GFP mouse cortex. The green color represents the lipids and hemoglobin signal, while the blue represents the unmixed protein signal. After segmenting out the capillaries, Figure 4B shows the cortical cells in blue and the capillaries in red. We topically applied Neurotrace (NT) 500/525 to the exposed cortex before sealing the cranial window to identify the pericytes within the imaged volume. Figure 4C shows the TPEF image of Tie2-GFP (cyan) and NT (green). Tie2-GFP was excited using two-color two-photon excitation of 800 nm and 1040 nm, and NT was excited using one-color two-photon excitation of 1040 nm. Both dyes were collected using a 525/40 nm bandpass filter and spectrally unmixed after collection. The first step in label-free identification was to stratify each cell identified in SRS imaging into one of two classes: capillary-lining and non-capillary-lining. This was achieved by determining if a cell had overlapping pixels with a capillary. A cell without overlapping pixels was classified as non-capillary-lining cell (color-coded yellow in Figure 4D). A cell that has overlapping pixels was classified as a capillary-lining cell and further stratified based on colocalization with the fluorescent labeling. The three classes of capillary-lining cells are endothelial cells (Tie2^+^, NT^−^), pericytes (Tie2^−^, NT^+^), unlabeled cells (Tie2^−^, NT^−^), color-coded blue, green, and cyan in Figure 4D, respectively. Unlabeled cells likely contain other cell types that can occupy capillary space including astrocytes, oligodendrocytes, and microglia cells. Representative 3D renderings of each cell type are shown in Figures 4E-4G. We further quantified the density of capillary-lining cells and non-capillary-lining cells. Figure 4H shows that the percentage of capillary-lining cells decreases with age. This is due to a decrease in capillary-lining cells rather than an increase in non-capillary-lining cells as shown in Supporting Figure 4.

**Figure 4.**
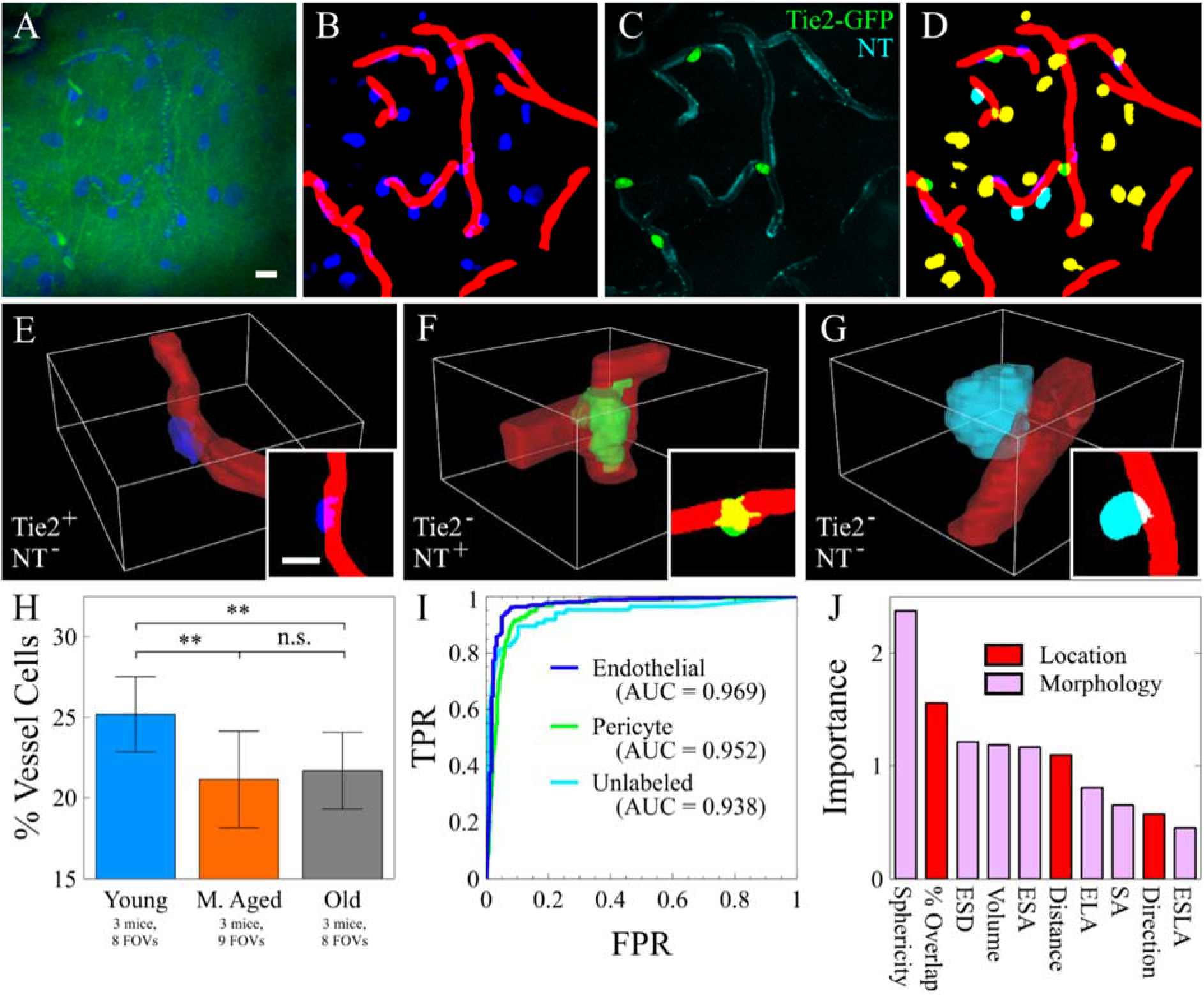
*In vivo* label-free cell identification of capillary-lining cells. A) SRS imaging intensity projection of 25 μm of Tie2-GFP mouse. Green shows SRS of lipids and TAM of hemoglobin. Blue shows the spectrally unmixed proteins. B) Spectrally unmixed proteins (blue) and capillaries (red). C) Two TPEF fluorophores: GFP (green) and Neurotrace 500/525 (cyan). D) Composite image of non-capillary-lining cells (yellow), unlabeled capillary-lining cells (cyan), pericytes (green), and endothelial cells (blue). E-G) 3D rendering of an endothelial cell, a pericyte, and an unlabeled cell, respectively, with corresponding 2D projection. H) Percentage of vessel cells out of total cell population with age. I) ROC curves of prediction accuracy for predicting endothelial cells, pericytes, and unlabeled cells based on morphological and locational features. J) Importance of each feature in the machine-learning classifier. Red features indicate the location of the cell to the capillary, pink features indicate morphology. Scale bars: 10 μm. **: p<0.01)

From here, we imaged nine Tie2-GFP mice ranging from P37 to P92 and extracted ten features from each capillary-lining cell we observed. Seven of the features were morphological: sphericity, equivalent spherical diameter (ESD), volume, ellipsoid short axis length (ESA), ellipsoid long axis length (ELA), surface area (SA), and ellipsoid second longest axis length (ESLA). The remaining three features were relational between the cell and neighboring capillary: percent volume overlap (% overlap), distance from cell centroid to capillary, and major axis angle between cell and capillary (θ ≤ 90°). These features were used to train a machine learning model for label-free cell prediction using SRS images alone. We achieved a 90 ± 2% prediction accuracy of 661 endothelial cells, 134 pericytes, and 120 unlabeled cells using a 20-tree Random Forest (RF) algorithm. The RF out-of-bag error plateaued near 20 trees, with negligible improvement from 20 trees to 100 trees (Supporting Figure 5). Figure 4I provides the ROC curves and area under the curve (AUC) for each cell type, and Figure 4J provides the relative importance of each feature in the RF algorithm. These results demonstrate the first *in vivo* cell identity prediction using SRS morphology. The SNARF method not only significantly improved the multiplexing capability of TPEF, but also enabled label-free imaging of specific cell types through machine learning. These capabilities will facilitate future studies of capillary function in normal aging and dysfunction in neurodegenerative diseases. Importantly, our label-free method can be potentially applied to any mouse which will allow researchers to study disease models without the requirement of dye application or transgenes.

## Discussion and Conclusion

The mouse brain is an extraordinarily complex organ with hundreds of different regions and over 100 million cells comprised of many different cell types.^57^ Understanding important biological processes of the brain such as neurovascular coupling, blood brain barrier, and neurodegeneration often require the ability to monitor many cell types and features simultaneously *in vivo*. TPEF has facilitated invaluable contributions to neurophotonics due to its high imaging resolution, versatility, and a wide variety of dyes and transgenic animals. However, despite its tremendous utility, TPEF still suffers from a few key limitations. First, TPEF spectra are typically broad (~50 nm), which can make detection of more than 3 fluorophores *in vivo* difficult. Spectral unmixing has recently been developed to address this challenge, but it still comes at the expense of system complexity and cost due to the use of multiple lasers and computational unmixing.^58^ Second, transgenic animals with multiple fluorescent proteins encoding different cell types require extensive breeding, which brings significant costs in time and resources. Third, while exogenous dyes can be used for labeling specific cells or structures, they also have limitations. Chemical dyes can alter the physiology of the brain. For instance, it has been reported that overexposure of sulforhodamine 101 (SR101), an astrocyte staining dye, can induce seizure-like brain activity in mice.^5^ Additionally, topical application and bolus injection for longitudinal studies become challenging to perform when using a sealed cranial window model. Intravenous injection mitigates the repeated loading problem but only affords a limited time window for imaging and has also shown unwanted side effects. For example, Hernandez *et al.* reported that intravenous application of a fluorescently labeled antibody used to stain WBCs within a stalled vessel lead to the clearing of stalled vessels 10 min after application.^52^ Although TPEF will continue to play a dominant role in high resolution *in vivo* brain imaging, these limitations highlight the need for alternative imaging methods, such as label-free multiphoton microscopy. The label-free multiphoton imaging techniques – stimulated Raman scattering (SRS) and transient absorption microscopy (TAM) – described in this study are orthogonal and complementary to TPEF. All three techniques are coupled into the same laser scanning microscope. They share the same excitation laser, which allows simultaneous detection of multiple contrasts. SNARF offers a richer characterization of brain structure and dynamics. The fluorescence imaging capabilities of SNARF can be further expanded to multiple fluorophores due to the use of two synchronized lasers at different wavelengths. We believe that SNARF multiplexing offers an exciting route to relieve some of the burdens of fluorescent labeling.

Multiphoton microscopy offers intrinsic optical sectioning, which facilitates straightforward morphological analysis. We leveraged this capability to quantify myelination in the cortex using SRS microscopy. Myelin dynamics play a crucial role in healthy brain development and homeostasis. As observed in normal aging and certain neurodegenerative diseases, demyelination occurs via the disruption of either the myelin sheath directly or the oligodendrocyte, the myelinating cell in the central nervous system. Identifying the cause of demyelination is essential for developing therapeutical treatments. As shown in this study, SRS can quantitatively imaging myelin and cell bodies simultaneously, which offers exciting potential to understanding myelin dynamics throughout normal aging and disease progression. We observed interesting and diverse balloon structures of the myelin sheath. They are more prevalent in middle-aged and old mice compared to young mice. These structures do not share the same features as observed with SCoRe, another label-free myelin sheath imaging technique. The implications of abnormal myelin sheath structure in aging and brain injury warrant further investigation of these structural changes using orthogonal techniques.

With the SNARF microscope, we demonstrated the first label-free TAM imaging of cortical hemodynamics. Hemodynamics is a key player in neurovascular coupling and overall homeostasis. We first investigated the recently reported phenomenon of stalled capillary flow in the cortex. With TAM, we observed three classes of blood vessels: flowing, transiently stalled (<15 min), and persistently stalled (>15 min). With SRS, we further quantified the presence of white blood cells (WBCs) within the persistently stalled vessels and found that ~45% (8/18) contained at least one WBC. Others do not have an apparent occlusion or capillary narrowing. Interestingly, we observed no change in the prevalence of stalled capillaries with age. Additional experiments are required to determine if the frequency of stalled WBCs changes in disease conditions. Because stalled flow can lead to local oxygen depletion and has been hypothesized to play a critical role in the development of neurodegenerative disease, it is important to explore the exact cause of different stalled flow conditions and oxygen delivery change with stalled flow. Although fluorescence tracers are convenient to visualize blood flow, we should exercise caution when using those tracers due to their potential in altering hemodynamics. Another potential benefit of label-free TAM imaging compared to fluorescent tracers is that it is possible to acquire oxygen saturation information based on the difference in absorption between oxy-hemoglobin and deoxy-hemoglobin.^25^ This approach could potentially increase the speed in measuring oxygen tension by three orders of magnitude compared to current oxygen probes^59^, thus enabling measurement of oxygen delivery change in response to both transient and persistent stalled flow.

Lastly, we used SNARF to quantify and identify the cells that line the capillaries and compose the BBB. With two-color SRS, all the cell bodies can be visualized, which is the basis for stimulated Raman histology of *ex vivo* tissue.^14^ We quantified the populations of non-capillary-lining cells and capillary-lining cells. We found that the density of non-capillary-lining cells remained constant with age while capillary-lining cells decreased from young mice (P37-P48) to middle-aged mice (P202-P246). We further built a Random Forest classifier to identify capillary-lining cells based on morphology and relationship to the capillary structure. By using ten features – seven morphological and three relational – we demonstrated 90% accuracy in using label-free SRS imaging to predict three classes of cells: endothelial cells, pericytes, and unlabeled capillary-lining cells. To our knowledge, this is the first report of using SRS to distinguish different cell types in tissues. We note that it is possible to extend the prediction to other cell types, including astrocytes, oligodendrocytes, and microglia cells, with no additional cost in imaging time or complexity. The only requirement is the separate machine learning training of specific types of cells using SNARF. The prediction accuracy can be improved with larger training datasets, high signal to noise (SNR), higher spatial resolution, and better algorithm. Both SNR and resolution can be improved through aberration correction.^60^ Deep learning can potentially replace machine learning to offer better performance in cell prediction.^61^ In combination with genetic labeling of different subtypes of neurons, SNARF significantly expands the features that can be monitored simultaneously and provides a powerful platform for *in vivo* imaging of the structure and function of the brain cortex.

The main limitation of SNARF is the shallow penetration depth of SRS and TAM. Although we have demonstrated the best imaging depth of up to 300 μm to date, we are still limited to cortical layer 2-3. In comparison, TPEF can routinely image at a depth of 500-600 μm due to the high sensitivity of fluorescence detection. There are several routes to improve the penetration depth. Aberration correction has been shown to improve the SNR of TPEF in the mouse cortex by 5-fold.^60^ It can potentially provide a larger benefit for SRS and TAM because of the large chromatic aberration besides spherical aberration due to the use of two widely separated wavelength near 800 nm and 1040 nm. Longer wavelength excitation is another strategy that can improve imaging depth for SRS and TPEF^18,62^, but it remains to be seen whether TAM signal size will be significantly affected. Because SRS and TAM use longer pulses than TPEF with less photodamage, it is also possible to use lower rep rate lasers to boost the signal further. Lastly, Raman labels are another possibility to increase SRS signal significantly. While a label is still required in this case, the potential advantage is that Raman labels have much greater capacity for multiplexing than fluorescence labels.^63^ We believe that these technical advancements will further improve SNARF microscope to enable multiplex imaging of a wide range of cells, structures, and functions for studying brain function and diseases *in vivo*.

## Acknowledgements

The authors would like to thank Dr. Andy Shih for meaningful conversations and guidance with cranial window surgery. The authors would also like to thank the Animal Use Training Services (AUTS) team for the donation of animal tissues that were necessary for the technical development of this project. In particular, the authors would like to thank Kathy Andrich, Erika French, and Francesca Perrotta for their help and training. This study was funded by NIH R35 GM133435 (D.F.), the Beckman Young Investigator Award (D.F.), and the Eli Lilly Award (D.F.).

## Notes

### Competing Interest Statement

The authors have declared no competing interest.

