## Supporting figures for "*In Vivo* Simultaneous Nonlinear Absorption Raman and Fluorescence (SNARF) Imaging of Mouse Brain Cortical Structures"

### SUPPORTING INFORMATION

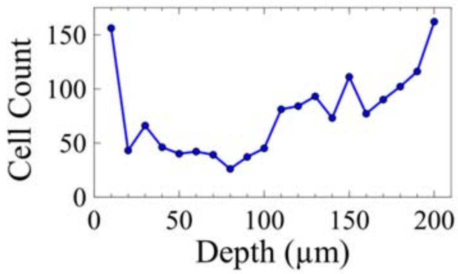


**Figure S1.** Depth dependent cell count for a P246 mouse. The increase in cells near 100 μm represents the transition from Layer 1 to Layer 2 of the cortex.

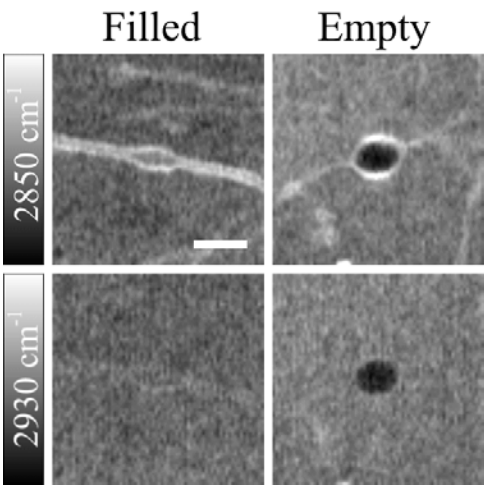


**Figure S2.** Contents of myelin balloons are either filled (left) or empty (right) observed in Layer 1 of a P246 mouse brain cortex. Scale bar: 5 μm.

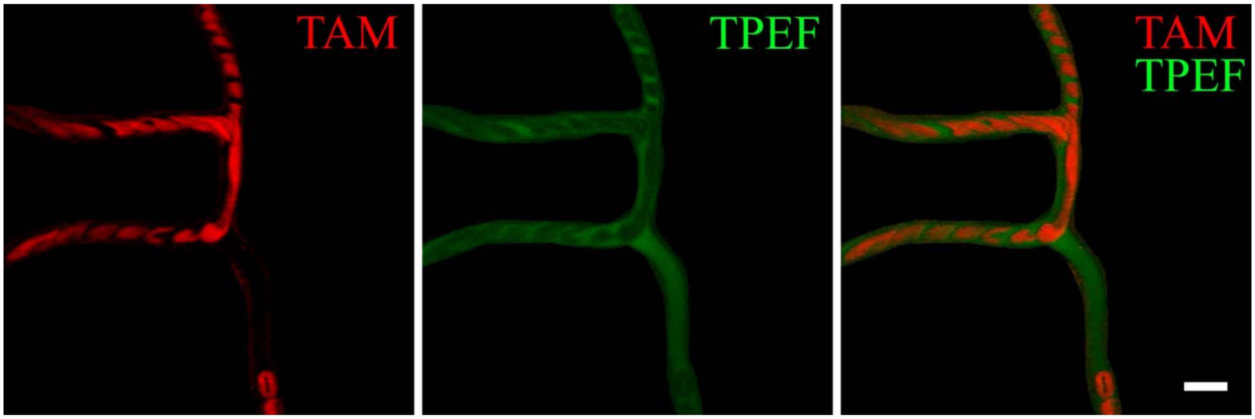


**Figure S3.** *In vivo* imaging of a mouse brain capillary using the intrinsic optical absorption of hemoglobin (red). The capillary is also stained with FITC (green) to demonstrate the orthogonality of the two techniques. Scale bar: 10 μm.

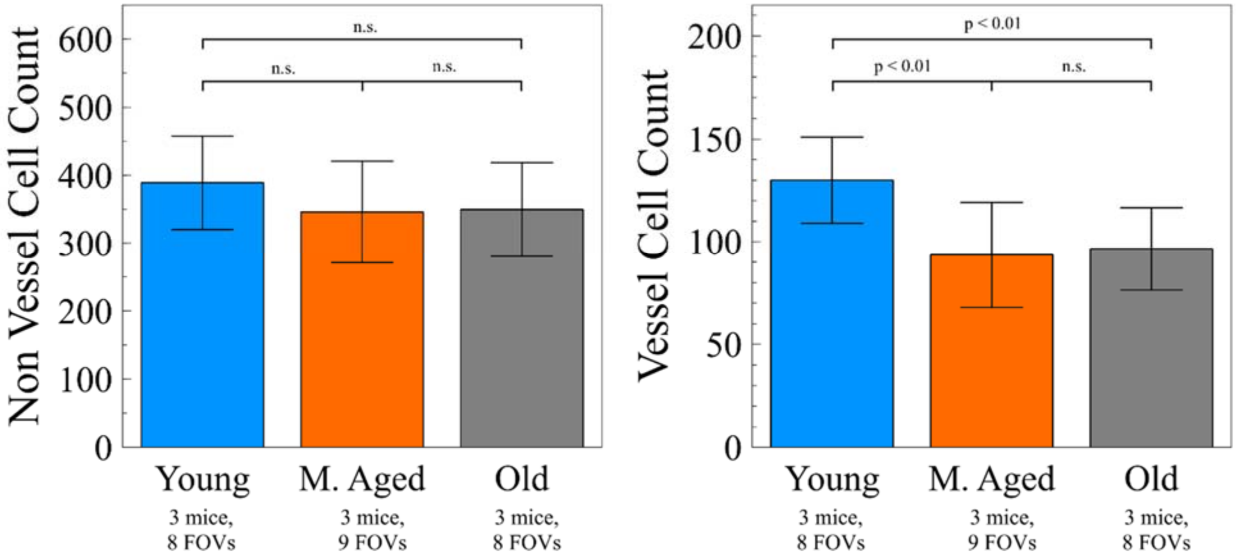


**Figure S4.** Cell populations in the top 200 μm as a function of age.

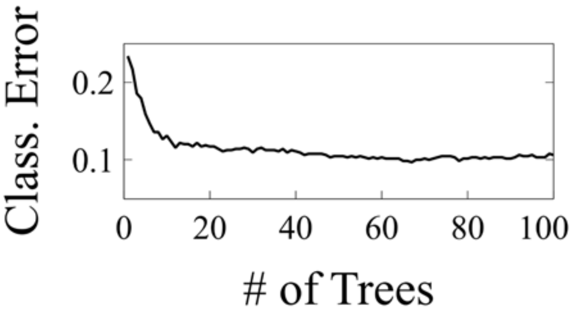


**Figure S5.** Prediction error as number of trees increases.
